# Differentiated genomic footprints and connections inferred from 440 Hmong-Mien genomes suggest their isolation and long-distance migration

**DOI:** 10.1101/2023.01.14.523079

**Authors:** Guanglin He, Jing Chen, Yan Liu, Rong Hu, Peixin Wang, Shuhan Duan, Qiuxia Sun, Renkuan Tang, Junbao Yang, Zhiyong Wang, Xiaofei Xu, Yuntao Sun, Libing Yun, Liping Hu, Jiangwei Yan, Shengjie Nie, Lanhai Wei, Chao Liu, Mengge Wang

## Abstract

**Background:** The underrepresentation of Hmong-Mien (HM) people in Asian genomic studies has hindered our comprehensive understanding of population history and human health. South China is an ethnolinguistically diverse region and indigenously settled by ethnolinguistically diverse HM, Austroasiatic (AA), Tai-Kadai (TK), Austronesian (AN), and Sino-Tibetan (ST) people, which is regarded as East Asia’s initial cradle of biodiversity. However, previous fragmented genetic studies have only presented a fraction of the landscape of genetic diversity in this region, especially the lack of haplotype-based genomic resources. The deep characterization of demographic history and natural-selection-relevant architecture in HM people was necessary.

**Results:** We comprehensively reported the population-specific genomic resources and explored the fine-scale genetic structure and adaptative features inferred from the high-density SNP data in 440 individuals from 34 ethnolinguistic populations, including previously unreported She. We identified solid genetic differentiation between inland (Miao/Yao) and coastal (She) southern Chinese HM people, and the latter obtained more gene flow from northern East Asians. Multiple admixture models further confirmed that extensive gene flow from surrounding ST, TK, and AN people entangled in forming the gene pool of coastal southeastern East Asian HM people. Population genetic findings of isolated shared unique ancestral components based on the sharing alleles and haplotypes deconstructed that HM people from Yungui Plateau carried the breadth of genomic diversity and previously unknown genetic features. We identified a direct and recent genetic connection between Chinese and Southeast Asian HM people as they shared the most extended IBD fragments, supporting the long-distance migration hypothesis. Uniparental phylogenetic topology and Network relationship reconstruction found ancient uniparental lineages in southwestern HM people. Finally, the population-specific biological adaptation study identified the shared and differentiated natural-selection signatures among inland and coastal HM people associated with physical features and immune function. The allele frequency spectrum (AFS) of clinical cancer susceptibility alleles and pharmacogenomic genes showed significant differences between HM and northern Chinese people.

**Conclusions:** Our extensive genetic evidence combined with the historic documents supported the view that ancient HM people originated in Yungui regions associated with ancient ‘Three-Miao tribes’ descended from the ancient Daxi-Qujialing-Shijiahe people. And then, some recently rapidly migrated to Southeast Asia, and some culturally dispersed eastward and mixed respectively with Southeast Asian indigenes, coastal Liangzhu-related ancient populations, and incoming southward Sino-Tibetan people. Generally, complex population migration, admixture, and adaptation history contributed to their specific patterns of non-coding or disease-related genetic variations.

## BACKGROUND

Comprehensive representative genomic resources from ethnolinguistically diverse populations can provide a complementary genetic landscape to understand who we are, how we get here, and why we are differentiated [1-4]. Geographically and ethnolinguistically high-coverage human genomic projects can also help us understand the genetic architecture of human disease and complex physical traits [4-8]. More and more genetic researchers and projects have identified that missing diversity in human genetic studies has hindered some medical applications and comprehensive understanding of genetic background in non-European populations as the existing European bias in medical and population genetic research [9]. Genetic analyses of worldwide people in the Human Genetic Diversity Project (HGDP) [10, 11], Simons Genetic Diversity Project (SGDP) [8], and the Estonian Biocentre Human Genome Diversity Panel (EGDP) [4] provide the basic patterns of human genetic diversity, admixture traces and migration models. Recently, regional genomic projects of GenomeAsia 100K Project [7], NyuWa genome resource, and China Metabolic Analytics Project (ChinaMAP) [5, 6] also further highlight the importance of identifying the missing genetic diversity, patterns of genetic admixture signatures and their medical relevance [7] in understudied human populations.

China has rich human biological diversity, and over six language families exist here, including Altaic (Mongolic, Tungusic, and Turkic), Sino-Tibetan (Sinitic and Tibeto-Burman, TB), Hmong-Mien (HM), Tai-Kadai (TK), Austronesian (AN), and Austroasiatic (AA). The genetic patterns of modern Chinese populations showed the population stratification among ethnolinguistically different people, which was strongly correlated with geography, culture, and language families [12-14]. Recent genetic cohorts from ChinaMAP [6] and NyuWa genome resources [5] have provided crucial genetic variation data from geographically different Chinese populations and offered new insights focused on population structure and the medical relevance of Chinese people. We also noticed that all these genetic studies in China mainly included Han Chinese as their major studied subjects, which would introduce the Han bias in Chinese population genetic studies and influence the health inequality of genomic benefit in the precision genomic medicine era. China had two independent agriculture innovation centers in the Yangtze River (millet) and Yellow River (rice) Basins. Rich civilization’s history of social development and technological innovation in the middle Holocene epoch facilitated the formation of the ancient Yangshao tribe and the Dawenkou tribe in North China and the Sanmiao tribe and Liangzhu society in South China. Recent ancient DNA has identified the genetic differentiation between northern and southern East Asians since the early Neolithic period and then experienced extensive population admixture events [15]. The patterns of evolutionary history in East Asia differed from those in Europe and Oceania, which had undergone large-scale population admixture and replacement [16]. Ancient human gene flow events out of East Asia have limited influence on the genetic backgrounds of East Asians [17]. However, ancient DNA from spatiotemporal East Asians has identified regional-restricted ancient founding lineages and contributed to the reconstruction of subsequent extensive population migration and admixture events [2, 3, 15, 18]. The significant bidirectional gene flow between the Yangtze River Basin rice farmers and Yellow River Basin millet farmers has significantly reshaped the patterns of genetic affinity and spectrum of allele frequency (AFS) among ethnolinguistically diverse East Asians [15, 18]. Ancient population substructures in East Asia were also observed in modern East Asians. Our previous genetic studies identified genetic substructures in the Amur River, Tibetan Plateau, and inland and coastal South China associated with geographic locations and linguistic affiliations [13, 19-21]. However, these efforts have been made primary foundational work to dissect the mystery of Chinese populations’ evolutionary and adaptive history. Fine-scale genetic structure of different people and patterns of genetic relationship and admixture of some Chinese populations should be further explored, especially in some ethnolinguistically diverse regions in South China.

Although ancient and modern autosomal genomic variations could construct complex population admixture models and unidentified human demographical events [2, 10, 20], whole-genome sequencing (WGS) of mitochondrial DNA and Y-chromosome could provide additional evolutionary traces based on the shared or novel haplotype groups, also referred to as the haplogroups. Wei et al. first used this approach to identify novel lineage informative markers among diverse genomes and constructed one recalibrated phylogenetic tree to explore population divergence and expansion events in the historic and prehistoric periods [22]. Poznik et al. analyzed 1,244 worldwide Y-chromosome sequences from the 1000 Genomes Project (1KGP) to characterize the landscape of Y-chromosome diversity of 26 populations. Karmin et al. investigated 456 geographically diverse high-coverage Y-chromosome sequences to construct the revised phylogenetic topology with the divergence time estimation of key mutation events [23, 24]. They have reported punctuated bursts and population bottlenecks associated with the cultural exchanges among worldwide continental populations [23, 24]. Regional Y-chromosome-based phylogenic investigation also reported shared ancestral or population-specific founding lineages and corresponding rapid expansions [25, 26]. Maternal lineages among different populations could also trace the process of population evolutionary past. Recent mitochondrial studies from modern and ancient Tibetan genomes have illuminated the Neolithic expansion processes of the Yellow River farmers and the Paleolithic peopling of the Tibetan Plateau [27-29]. Li et al. also reported that the maternal structure of Han Chinese was stratified along three main Chinese river boundaries [30]. It is obvious that fine-scale and large-scale uniparental genetic studies should be conducted to explore the evolutionary history of the understudied Chinese ethnolinguistic populations [31, 32].

HM-speaking populations include those who speak Hmong (Miao and Hmong) and Mien (Yao, She, and Dao) in the Yungui Plateau and Wuyi mountainous areas of South China, Vietnam, and Thailand in northern Southeast Asia [33]. The original homeland of HM people was suggested to be in Central China, associated with Neolithic Shijiahe, Qujialing, and Daxi cultures in northern regions of Yungui Plateau. Historic documents showed that exiling from Central China by Han Chinese expansion or other ancient northern East Asians promoted the southward of ancient HM people [34]. The complex migration and admixture history needed to be further explored. The interaction between HM-speaking people and other southern Chinese populations (TK, ST, AN, and AA) also required further exploration. Recent findings from the uniparental mitochondrial DNA, Y-chromosome, and genome-wide evidence have identified different evolutionary processes between inland TK and HM people and coastal AN and TK people [35]. Wen et al. found some HM-specific maternal lineages (B, R9, N9a, and M), and northern East Asian dominant lineages suggested extensive maternal interaction between northern and southern East Asians [36]. Wang et al. recently also identified the extensive population admixture between HM people and Chongqing Han [37]. Interestingly, a similar pattern of the unique and differentiated genetic structure of Sichuan Miao (SCM) has been reported by Liu et al. [38]. They identified the close genetic relationship between SCM and Vietnam HM people. Recently, pilot WGS work conducted by Xu et al. has reported the evolutionary history of HM-speaking Yao in Guangxi and suggested that Yao diverged from the other southern Chinese in the middle Neolithic period (9700 YBP) and separated within HM people at 6700 YBP. However, the comprehensive landscape of admixture and migration patterns of geographically diverse HM people and the genetic resources of another important HM-speaking She keep unreported. She, with a census population size of 746,385, is widely distributed in central and southern China and one of the important parts of HM people, but around 90% of She people reside in Fujian and Zhejiang. Historians proposed different originated hypotheses about the She people, including the Wuliang Man, Dongyi, and Yue people’s decedents and Nanman and Min’s decedents. However, no genetic studies have been conducted to resolve this problem. Thus, we collected genome-wide data from all HM-speaking ethnic people, including newly-collected coastal HM people, and conducted a comprehensive population genetic analysis based on sharing alleles and haplotypes to explore the following key questions: 1) what’s the landscape of genetic admixture and evolution within and between HM people; 2) How about the genetic relationship between HM people and other modern and ancient East Asians; 3) How about the connection among geographically distant HM people; 4) How about the evolutionary history of HM people and their interaction with surrounding neighbors? Our findings presented the full landscape of variation profiles, admixture process, biological adaptative history, and medical relevance based on the HM genomic resources.

## Results

### General population admixture patterns of HM people in the context of modern and ancient East Asians

To comprehensively explore genetic diversity and illuminate the evolutionary relationship among geographically diverse HM genomes, we generated high-density SNP data in 349 individuals from 26 Chinese populations (**Figure 1a∼b**). Previously published 20 HM people (She and Miao) from HGDP and 71 HM individuals from Vietnam and Thailand reported by Stoneking et al. were also included here to explore the genetic heterogeneity and connection among geographically diverse groups [39-41]. We obtained the final HM dataset, including 440 individuals from 34 populations, to formally investigate the evolutionary and adaptative history of HM-speaking people from South China and Southeast Asia. We also merged our data with publicly available 929 high-coverage WGS genomes from 53 worldwide populations included in the HGDP and 317 high-coverage WGS genomes from 26 populations in Oceanian genomic resources, 20580 unique modern and ancient individual genotype data included in the Human Origin (HO) and 1240K datasets from the Allen Ancient DNA Resource (AADR) [39, 42, 43], and the merged high-density Illumina or WGS dataset, the merged middle-density 1240K dataset and the merged low-density HO dataset were generated respectively (**Table S1**). The former one was mainly used for the identification of natural-selection signatures, and the latter two datasets were used primarily for population admixture modeling and demographic history reconstruction.

**Figure 1.**
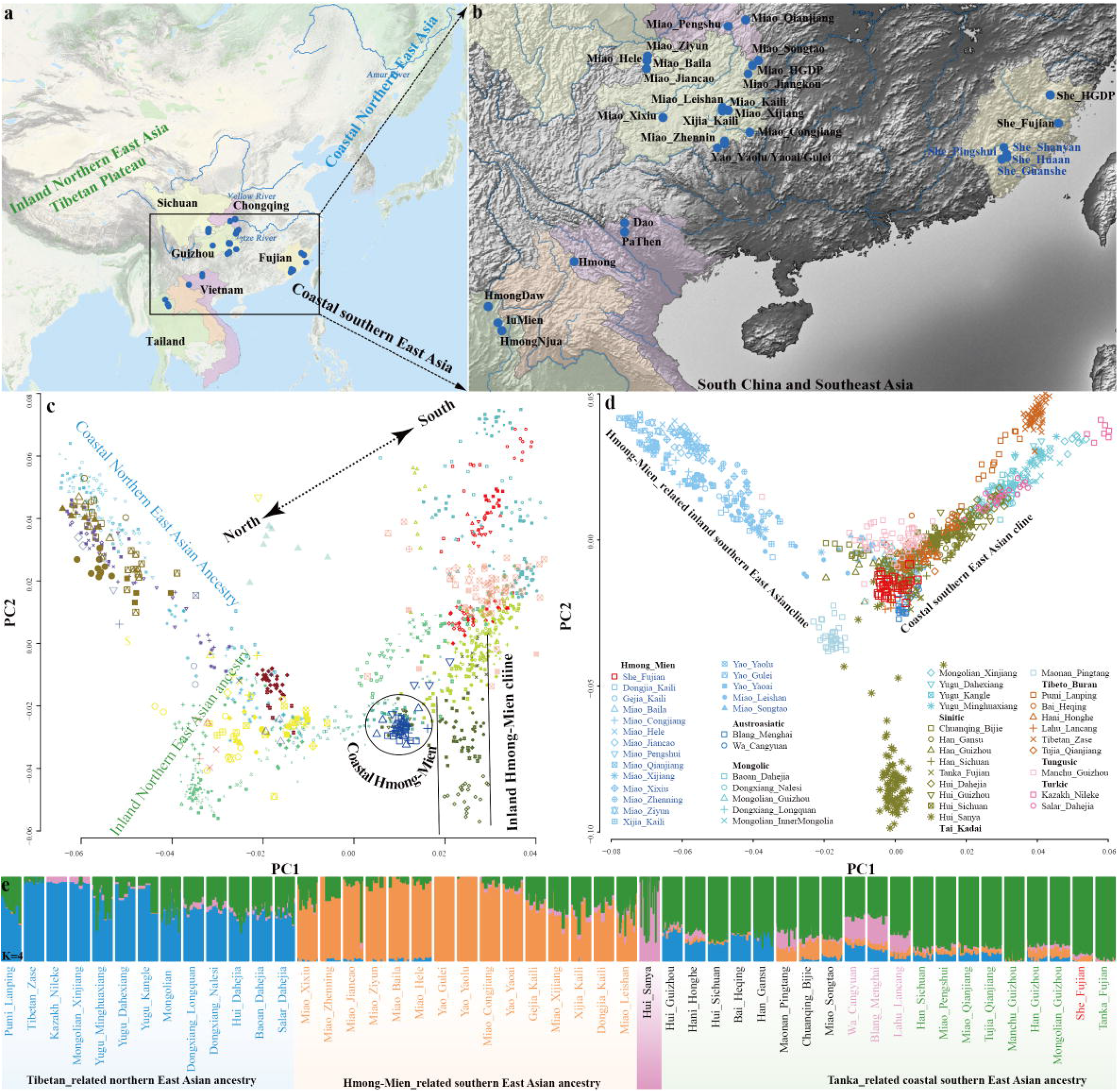
Geographical positions and genetic admixture of Hmong-Mien (HM) people. (**a∼b**) The map of Eastern Eurasia included China and Southeast Asia. Regions of all included HM population collection were zoomed in right regional map of South China and Southeast Asia. (**c**) Principal component analysis showed the general patterns of the genetic background of modern and ancient East Asian based on the merged Human Origin dataset, where ancient people were projected on the PCA. Modern populations from different language families and ancient populations from different geographical regions were color-coded. The detailed legends for Figure 1c are presented in **Figure S1**. (**d**) PCA showed the genetic patterns of Chinese populations based on the high-density Illumina dataset. (**e**) Model-based ADMIXTURE results with four predefined ancestral sources showed the individual or population composition.

To investigate and present the basic pattern of the genetic admixture and the relationship between HM people and other modern and ancient populations, we conducted principal component analysis (PCA) and unsupervised ADMIXTURE among modern and ancient Eastern Eurasians. We projected the ancient people into modern population backgrounds (**Figure S1∼2**). Clustering patterns based on the genetic variations among Siberian and East Asian peoples showed significant differentiation between ancient southern Siberians, Northern East Asians, and Southern East Asians (**Figure S1a**). We further conducted PCA tests focused on people from Southeast Asia and East Asia to explore the fine-scale relationship among the regional populations (**Figure S1b**). Fine-scale population substructures were dissected and respectively associated with geographical regions and cultural features: Mongolic and Tungusic people from Northeastern China clustered with ancient Neolithic Boisman, DevilsCave, and Baikal Lake people. TB-speaking Tibetan people clustered together and grouped closely to the Nepal Chokhopani, Mebrak, and Samdzong people (**Figure 1c)**. Southern East Asians were also separated into two major groups. Coastal AN people formed a genetic cline and clustered with Neolithic Fujian and Taiwan Hanben people. Inland HM-speaking Hmong, Dao, and PaThen formed a cline. However, She and Miao from HGDP huddled closely with Han Chinese, suggesting the stratification among HM people. We could confirm the identified population stratifications within the regional populations from inland and coastal South China and Southeast Asia (**Figures S1c**). We also explored the patterns of genetic substructure among Chinese populations using the merged high-density SNP data and found clustering patterns and ancestral components from HM people were significantly different from other groups (**Figures 1d∼e**).

The ancestry composition of 254 modern and ancient people inferred from the model-based ADMIXTURE presented the significant genetic differentiation between northern and southern East Asians as observed in the PCs. We observed four ancestral components maximized in northern East Asians related to Lubrak, Mongolia_N_North, Tarim_EMBA1, and Japan_Jomon, and four ancestral sources in southern East Asians and Southeast Asians related to Htin_Mal, Mlabri, Taiwan_Hanben, and Yao_Gulei (**Figure 2a**). Ancient Gaohuahua people from Guangxi and modern HM-speaking Yao, Miao, Hmong, PaThen, and IuMien possessed a similar ancestral composition and had the highest proportion of bright green ancestry. To confirm the identified pattern of genetic diversity, we excluded some reference populations in our dataset and conducted additional ADMIXTURE analyses (**Figures S1∼2**). We observed the consistent patterns of ancestry composition and found that four ancestry components enriched in AA-speaking Htin, HM-speaking Hmong, AN-related Hanben and Ami, and northern ancestry related to Neolithic Shamanka people when we predefined four ancestral sources. When ancestral sources increased to five, inland northern East Asian Tibetan-related ancestry was separated from others (**Figures S1∼2**). Inland HM people showed their specific ancestral component. The genetic structure of the coastal She people from Fujian was first reported here and showed a genetic ancestry similar to geographically close populations, not to the linguistically close inland HM people. Admixture scenarios among 34 HM groups were further explored using the two clustering approaches implemented in ADMIXTURE and fineSTRUCTURE (**Figure 2b∼c**). The resulting patterns of genetic substructures inferred from the two methods were consistent, reflecting the genetic affinity among geographically close populations (**Figure S3**). Generally, population structure inferred from the PCA and ADMIXTURE suggested that complex population separation and admixture played a pivotal role in the basic landscape of modern and ancient East Asians. Geographically different HM people harbored complex and differentiated population history. Based on the observed patterns of genetic differentiation among geographically diverse HM populations, we followingly comprehensively characterized the genetic history of coastal HM She. Then we explored the whole landscape of the demographical history and medical relevance of all geographically diverse HM people based on the state-of-art methods in the following sections.

**Figure 2.**
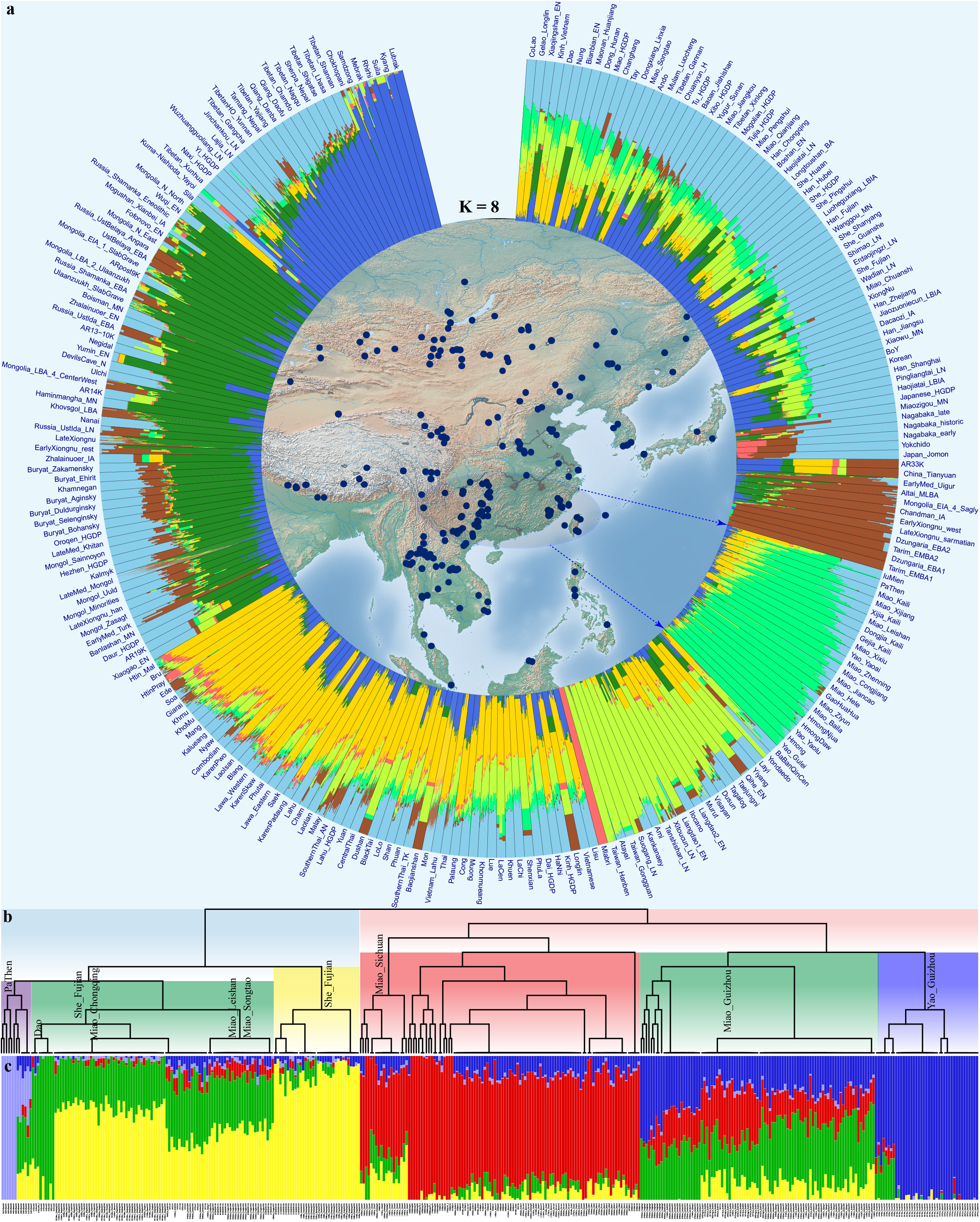
Population admixture and genetic ancestry among 153 modern ethnolinguistically diverse eastern Eurasians and 101 ancient populations from East Asia and surrounding regions. (**a**) The ancestral composition of 254 modern and ancient populations was inferred based on the genetic variations in the merged Human Origin dataset. The error of the cross-validation showed the model with eight ancestral sources was the best-fitted model. The distribution of all included populations was shown in the central Map. (**b**) Fine-scale population structure among 34 HM people inferred from the shared haplotypes and (**c**) the ancestral composition with five predefined ancestral sources among them inferred based on the allele-sharing patterns.

### Demographical history reconstruction of southeastern coastal HM people

Genome-wide population variations and patterns of genetic diversity of three HM-speaking She populations and one AN-speaking Gaoshan group from Fujian in southeastern coastal China were first reported here, where the Gaoshan people were used as the control for exploring the similarity between HM and mainland AN people. We studied genetic admixture patterns and the ancestry composition of She people in the context of 175 modern and ancient East Asians and Southeast Asians using model-based ADMIXTURE and PCA. We hypothesized the number of ancestral sources ranging from 2 to 11 and observed differentiated genetic features of She people in different dissection resolutions (**Figure S2**). Hmong-related ancestry in She people (including one She population from HGDP) ranged from 0.902 to 0.910. Northern East Asian ancestry related to Tibetan and Siberian people were separated from each other when K=3∼5. She people derived ancestry from Tibetan-related and Hanben-related people. When the ancestry sources were assumed to be 6∼8, southern East Asian ancestries related to HM and AN people were separated. We found that the She people were modeled with primary ancestry from Hmong and Tibetan and minor ancestry from AN people. Interestingly, when K values were larger than 9, we identified one blue ancestry maximized in She people.

To localize the genetic relationship between modern and ancient East Asians and explore the genetic relationship between She and geographically close ancient populations, we conducted PCA analysis among 1466 individuals from 179 populations. She and Gaoshan people were clustered with southern East Asians and had the most intimate relationship with Fujian ancient people (**Figure 1c**). When we excluded the ancient populations and only focused on the modern human populations, the She people showed a close relationship with Fujian Han Chinese and separated from the HM cline (**Figure S1**). Fine-scale population structure among southern Chinese and Southeast Asians showed that geographically different She people clustered together and separated with Gaoshan people and HM people from Southeast Asia. The estimated pairwise Fst values between the four She people also showed that they had the closest genetic relationship with each other and Han Chinese populations from South China, followed by southern Chinese TK people (**Table S2**).

To explore the genetic affinity between Coastal HM and Gaoshan people and modern and ancient East Asians, we calculated the outgroup-*f*_*3*_(Eurasian, She/Gaoshan; Mbuti). The estimated genetic drift showed that She people shared the closest genetic relationship with geographically close She people and Han Chinese from Fujian and Guangzhou provinces in modern reference datasets and Bronze Age to historical people from Henan province (Luoheguxiang and Pingliangtai, **Table S3**). Three population tests in the form *f*_*3*_(source1, source2; targeted populations) could provide direct evidence for genetic admixture signatures with statistically significant Z-scores (Z<-3). However, we found no admixture signatures focused on the targets of the She people. We identified many negative values in *f*_*3*_(Ami, Eurasian; Gaoshan), which suggested that the Fujian Gaoshan people were one mixed group with one source from Ami and another associated with Han Chinese (**Table S4**).

To test which populations shared the most alleles with our surveyed She and Gaoshan people, we conducted formal *f*_*4*_-statistics in the form *f*_*4*_(reference population1, reference population2; studied She/Gaoshan, Mbuti). Pingshui She shared most alleles with She_HGDP compared with our used inland HM and other Asian reference populations except for Shanghai Han, Haojiatai_LN, and Longtoushan_BA (**Figure S1**). Similar patterns of shared ancestry between She and Han Chinese populations were also identified in She_Shanyang and She_Guanshe (**Figures S4∼7**). Different from the status of the She people, Gaoshan people shared the most alleles with AN-speaking Ami and Kankanaey compared with other reference populations, followed by TK-speaking people and Han Chinese (**Figure S4∼7**).

Modern and ancient northern East Asians shared more alleles with Fujian Gaoshan people compared to reference populations from other AN, AA, and TK people in South China and Southeast Asia. It was suggested that Gaoshan people shared more northern East Asian ancestry compared with ANs and other southern East Asians. Similarly, the values of the estimated statistically negative *f*_*4*_(ST/ancient northern East Asians, Gaoshan; AN/TK/AA, Mbuti) suggested that Fujian Gaoshan people harbored more ancestry related to southern East Asians related to AN (**Figure S7**). Strong shared ancestry derived from AN people in Fujian Gaoshan people was further confirmed in the negative values in the *f*_*4*_(Eurasian reference populations, Gaoshan; Ami/Kankanaey/Hanben, Mbuti). Results from the negative values in *f*_*4*_(Ami/Kankanaey/Hanben, Gaoshan; northern East Asian reference populations, Mbuti) further confirmed that Fujian Gaoshan people obtained genetic influence related to northern East Asians compared with Taiwan island populations.

Formal tests in *f*_*4*_(Han_Fujian/She_HGDP, She_Guanshe; Eurasian reference populations, Mbuti) have not identified statistically significant values, suggesting that studied She people formed one clade with Han Chinese and She_HGDP. Next, we tested which population may be the ancestral source of the She people. We hypothesized that She people were directly derived from northern East Asians and then tested *f*_*4*_(Northern East Asians, She; Eurasian reference populations, Mbuti). However, we identified many statistically negative values when we used populations from South China and Southeast Asia as ancestral source proximity. Our identified signals suggested that She people obtained additional ancestry from southern East Asians. Similarly, compared with southern East Asians, She people also obtained additional ancestry from northern East Asians. Taking it together, we could conclude that coastal HM was genetically close to geographically close Han Chinese and harbored ancestries from northern and southern East Asians. To further dissect the different ancestral origin hypotheses of She people, we conducted *f*_*4*_(Dong/Li/Zhuang/Maonan/Mulam/Gelao/inland HM, She; Eurasians, Mbuti) and identified many negative values when we used ST and AN people as the reference populations. We confirmed our results by replacing She with the other two studied populations (**Figure S8∼11**).

To explore the possible demographical models for illuminating these populations’ admixture models and admixture times, we used different statistical methods and ancestral sources to examine the admixture sources, proportions, and times. We first used geographically close Iron Age Hanben people from Taiwan as the southern East Asian sources, used ancient millet farmers from the Yellow River Basin as the northern sources, and conducted the qpAdm to test the two-way admixture models for She and Gaoshan. We found that all four studied populations were modeled as primary ancestry from Hanben people and minor ancestry derived from Yangshao or Longshan people. We reconstructed the qpGraph-based topologies with different northern and southern East Asian lineages (**Figure 3a**). We found two-way admixture models fitting all coastal HM populations, in which Gaoshan and She from Pingshui harbored more ancestry related to the Hanben people. We further dated the admixture events based on the ancestral sources from northern and southern East Asians. We found that She people harbored the admixture signatures since four thousand years ago with the ancestry contamination from northern and southern Chinese, which was further confirmed with Globetrotter results (**Table S5, Figure 3b∼c**).

**Figure 3.**
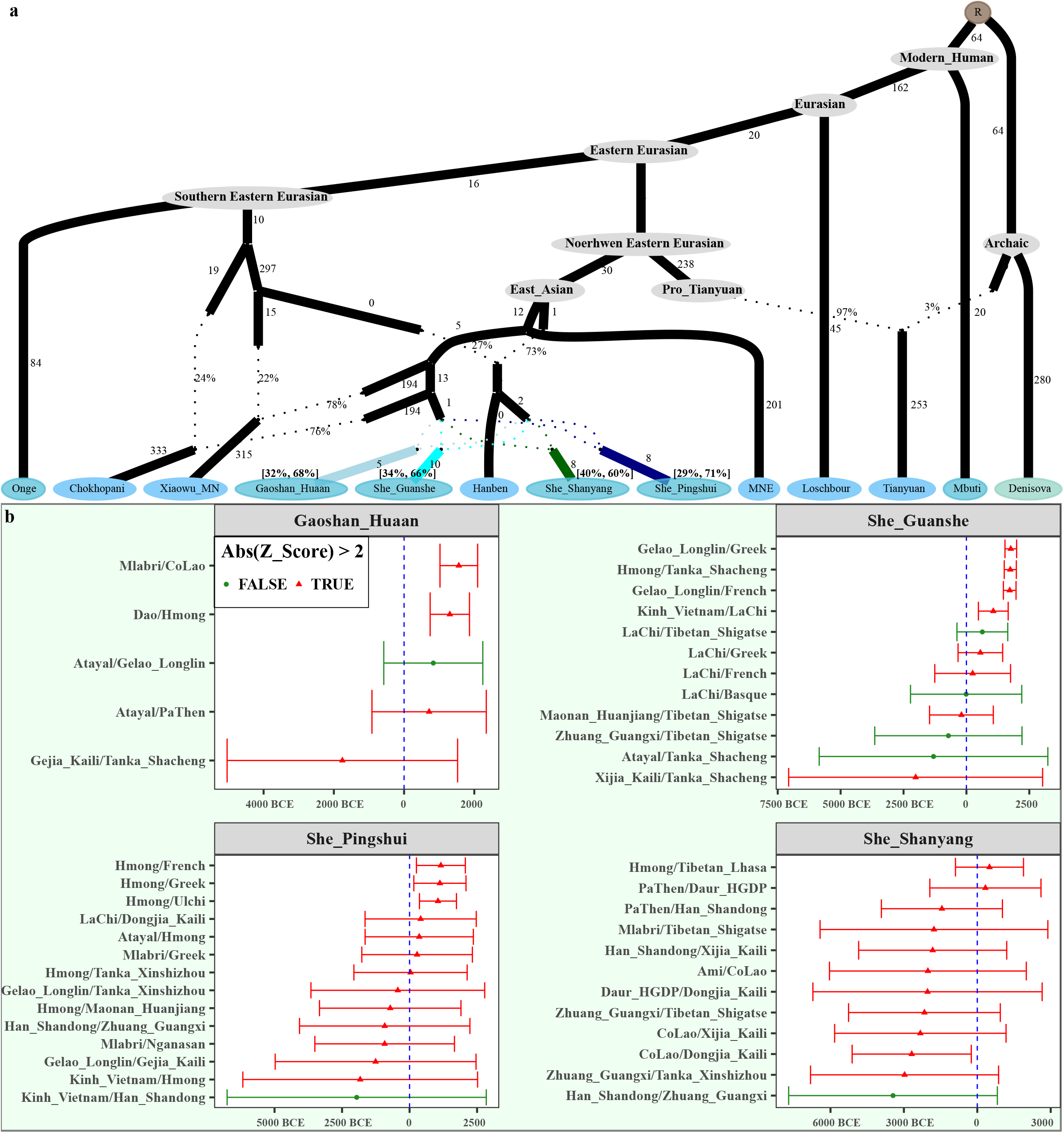
Demographic history of southern coastal HM people. (**a**) qpGraph model showed the evolutionary history of HM and corresponding ancient East Asian lineages. The presented qpGraph-based phylogenetic topology was fitted for Gaoshan people with the best-worst Z-score of −1.992. We manually added to admixture events of She people in this model. The Z-scores of the Guanshe people are 1.878, 1.873 for Pingshui She and −2.258 for Shanyang She. Branch lengths were labeled with 1000 times genetic drift. The dotted line indicates the admixture events with admixture proportion. (**b**) Admixture time estimates using different ancestral sources.

### Fine-scale population differentiation between inland and coastal HM-speaking populations

We comprehensively explored the genetic structure among 349 unrelated individuals from 26 southern Chinese HM populations based on the merged Illumina dataset to reconstruct the fine-scale population demographical history of HM people. The constructed phylogenetic topology among 203 male individuals revealed that O1 and O2 lineages contributed the most gene materials to the HM people, which were regarded as the two major ancestral founder lineages of HM people (**Figure 4a**). Northern East Asian dominant lineages (C2a/b, D1a1a, Q1a2a, and N1b) related to Mongolian or Tibetan were also identified in HM people, suggesting that the consistent southward gene flow influenced the gene pool of our studied populations. We also constructed the phylogenic relationship using the high-density Y-chromosome SNPs to explore the pattern of population divergence and expansion (**Figure 4b**). Phylogenetic topology showed extensive genetic interaction among inland HM Miao and Yao, who were extensively assigned to different paternal lineages. We found that two significant lineages of O2a2a1a2a1a1 and O1a1a1a1a1a1 experienced recent expansion among HM people. O2a2a1a2a1a1 lineages were mainly observed in Yao and geographically close inland Miao people, and O1a1a1a1a1a1 lineages were mainly identified in She people. The different founder lineages observed among inland and coastal HM people supported the observed genetic differentiation in the PCA patterns.

**Figure 4.**
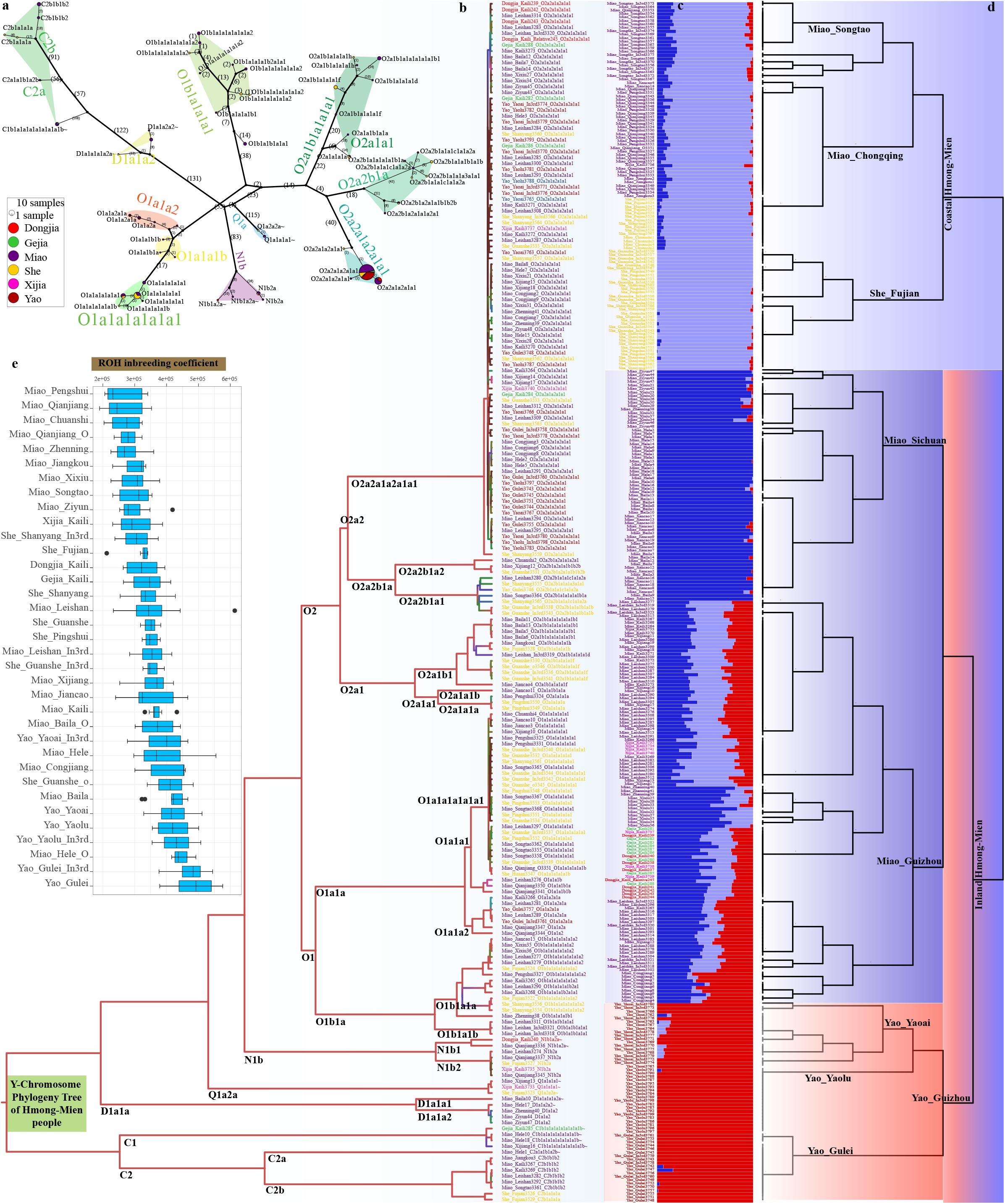
Fine-scale genetic structure of HM people inferred from genome-wide SNP data. (**a**) Median Joining Network showed the paternal lineages reconstructed via popART. (**b**) Individual lineages clustering patterns inferred from the LineageTracker. (**c**) Model-based ADMIXTURE with three predefined ancestral sources. The lowest cross-validation error value was observed when K=3. (**d**) The fineSTRUCTURE-based dendrogram showed a consistent clustered pattern with ADMIXTURE results. (**e**) The box plot showed the runs of homozygosity (ROH) of HM people.

To explore the fine-scale population genetic structures among 26 Chinese HM populations via the sharing alleles and haplotypes, we phased the 349 HM genomes and painted them as mosaic patterns using ChromoPainter. We explored the genetic similarities and differences using fineSTRUCTURE based on the sharing haplotype chunks (**Figure 4c**). The reconstructed dendrogram separated all included people into two major clades and several subclades. The upper clade included people from She and Chongqing Miao (CQM) people. Previous genome-wide SNP data revealed that the observed genetic diversity in CQM showed a similar pattern with geographical Han Chinese [37]. The acknowledged genetic affinity between CQM people and Coastal HM showed that both populations obtained significant genetic influence from Han Chinese populations. The lower clade included three subclades of people from SCM, Guizhou Miao, and Guizhou Yao people. Interestingly, we found Guizhou Yao separated from the other two Miao subclades first, and then the Guizhou Miao separated from the SCM subclade. We further conducted ADMIXTURE to dissect the ancestral composition based on the sharing AFS. Population substructures were also identified between geographically different HM people with the best-fitted models with three ancestral sources, suggesting the differentiated evolutionary processes within HM people. Coastal HM and CQM people had the maximum light-blue ancestry component. SCM possessed the highest deep-blue ancestry component, and Guizhou Yao people were dominated by the red ancestry component. Guizhou Miao people had three ancestries, as mentioned above. The reconstructed IBD-based phylogenetic topologies were strongly correlated with the population stratification as observed in model-based ADMIXTURE results (**Figure 4c∼d**).

The observed pattern of genetic structure among Chinese populations suggested that population isolation and admixture contributed to the formation of the gene pool of HM people. To explore whether other possible population evolutionary forces contributed to the complex pattern of genetic diversity, we estimated the Runs of Homozygosity (ROH) among these Miao, Yao, and She populations. We found the longest ROH indexes in Yao people than that in other populations (**Figure 4e**). Miao people have recently mixed with Han Chinese population and tended to have the shortest ROH values, which was consistent with previously evidenced admixture processes via admixture model reconstruction [37]. We also estimated the effective population size among HM people (**Figure 5a**), and all studied populations experienced different tracts of population bottleneck followed by population expansion. We also obtained the complex population interaction as evidenced via the estimated admixture signatures in the admixture three population tests and the identified differentiated shared signatures in the four population comparisons. Our results suggested that population admixture, separation, and interbreeding contributed to the genetic differentiation among HM people.

**Figure 5.**
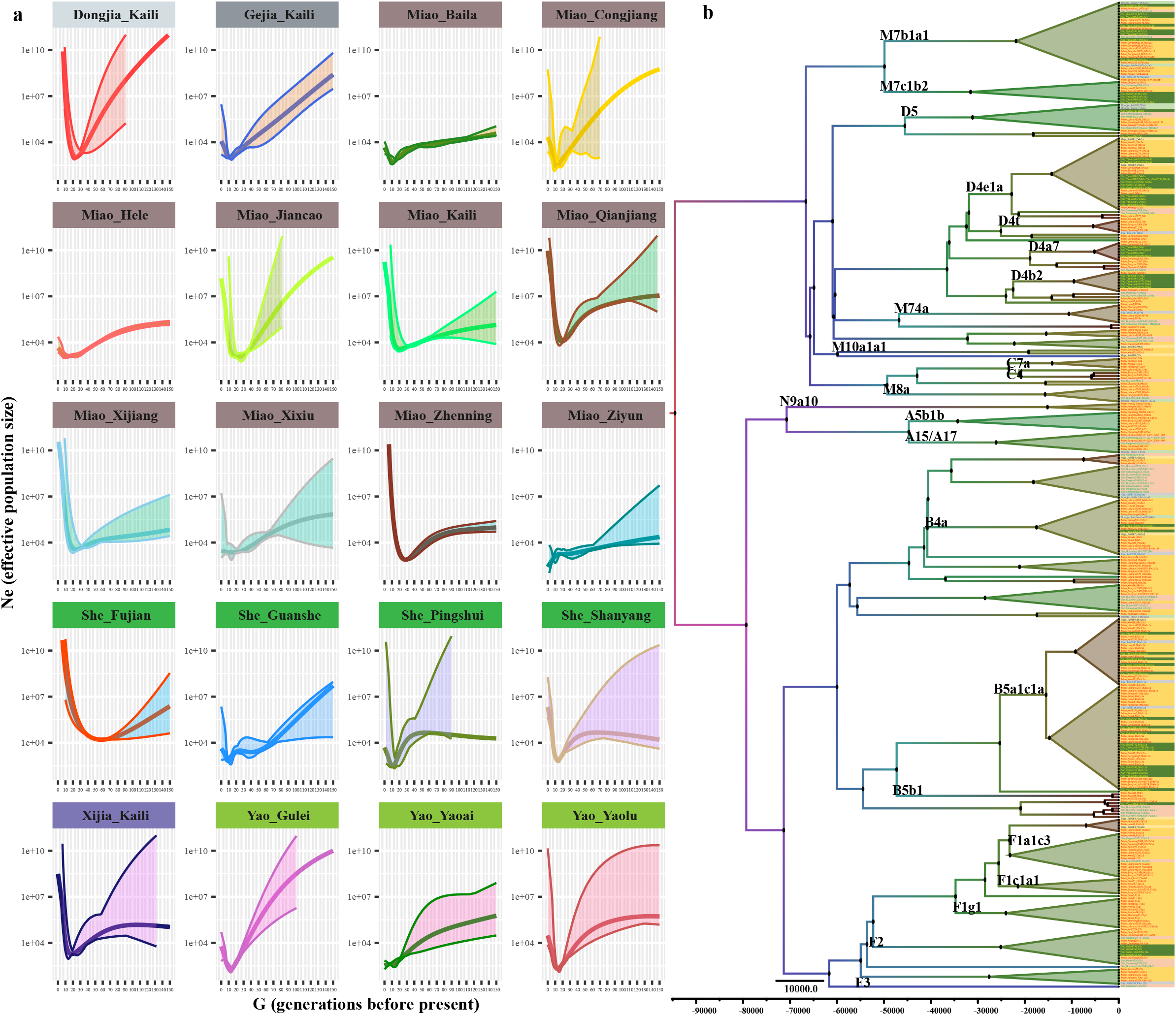
Effective population size and maternal phylogenetic tree of HM people. (**a**) The effective population sizes with the 95% confidence interval of HM people were estimated based on the IBD lengths. (**b**) The constructed mtDNA phylogeny showed the population divergence, expansion, and admixture among geographically different HM people.

Although the patrilocality of East Asians was supported by previous cultural and genetic evidence, we also explored the haplotype frequency spectrum (HFS) distribution, maternal phylogenetic topology, and Network-based phylogenetic relationship to study the maternal genetic structure of HM people. We observed that F, B, A, M, and N lineages contributed to the maternal gene pool of HM people. Based on the population size of defined terminal lineages, B5a1c1a, F1a1c3, D4e1a, and M7b1a1 were the founding lineages of Chinese HM people. The mosaic patterns of the ethnicity of one targeted lineage showed frequent maternal movement among ethnolinguistically diverse HM people (**Figure 5b**). The population movement and admixture patterns were also identified in the Network analysis. The identified founding lineages were distributed among different ethnic groups (**Figure 6**). We also identified star-like expansion in some lineages (D4, B5, and M7), supporting the recent population expansions.

**Figure 6.**
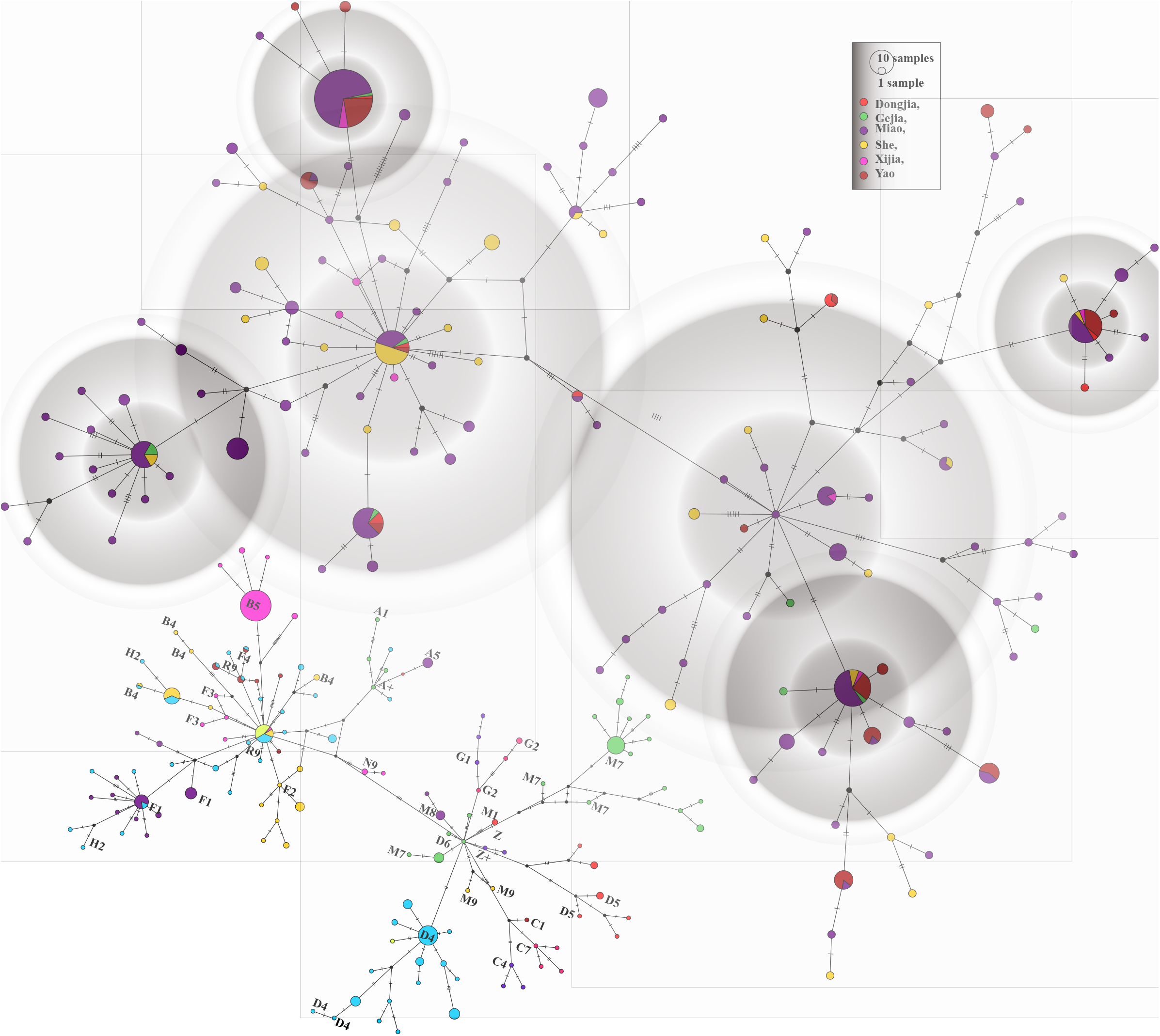
The network-based phylogenetic relationship showed the maternal genetic history of HM people. Some star-like population events were emphasized using the gray color. The haplogroup distribution was shown in the left bottom network topology.

### The direct genetic connection between inland HM and Southeast Asian HM people suggests the recent long-distance migration

Previous work has suggested that genes could be used to trace the common recent demographic history and recent human global population movements that have contributed to the complex admixture of previously geographically or culturally isolated populations, such as the Bantu expansion, the transatlantic slave trade, Mongolia empire expansion and Austronesian dispersal to Oceanian. ADMIXTURE and PCA results showed a close genetic relationship between inland HM and Southeast Asian HM people, which was also confirmed via the genetic distances (Fst and outgroup-*f*_*3*_ values). Ancient Gaohuahua people from Guangxi also had a strong genetic affinity with HM people from Yungui Plateau and Southeast Asia. To further explore the genetic connection based on the sharing haplotypes and visualize the affinity with geographical position, we explored the genetic connection among Chinese populations based on the high-density genetic markers and all HM populations from South China and Mainland Southeast Asia based on the merged 1240K low-density SNP data (**Figure 7**). The estimated IBD in different categories showed the extent of the population interaction in different periods. Among Chinese people, we identified extensive population interactions in different historic periods among geographically diverse inland Miao and Yao. Still, we only identified an ancient connection between the coastal and the inland HM people. Focused on all included HM people, we identified the recent genetic relationship between inland HM people from China and Mainland Southeast Asia, consistent with the historical documents of the long-distance migrations of HM people.

**Figure 7.**
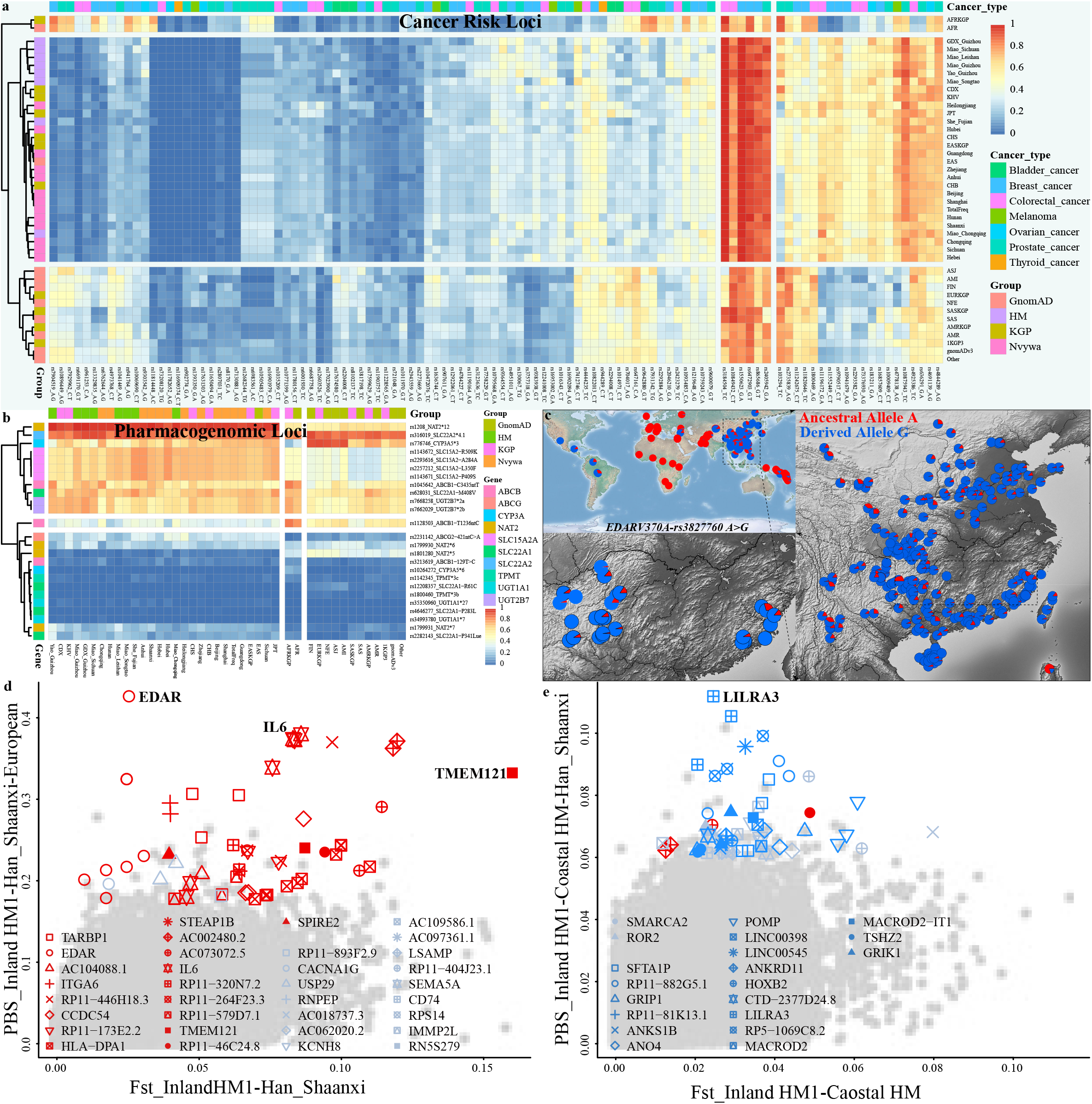
Genetic connection inferred from different IBD categories. (**a∼f**) The genetic connection among Chinese Miao, Yao, and She people showed their interaction status based on the high-density Illumina dataset. (**g∼j**) Shared IBD among all included HM people was estimated using the low-density merged Human Origin dataset.

### Medical relevance and local biological adaptation

Identification of risk alleles with small or large effect sizes in the cancer susceptibility genes via genome-wide association studies (GWAS) significantly influenced cancer clinical utility and epidemiological studies [44]. Population-based cancer prevention and screening programs depend highly on the risk discrimination estimated from the combined effect of multiple risk SNPs or individual high-effect inherited genetic susceptibility loci. To explore the AFS among our HM genomes and worldwide reference populations from NyuWa, 1KGP, and GnomAD, we investigated the genetic variations of 255 risk SNPs located in 180 genes from 27 different malignancies. The top included cancers were breast, prostate, colorectal, chronic_lymphocytic_leukemia diseases, and the top five gene annotation locations were intron, intergenic, missense, and upstream/downstream genes. The calculated AFS of risk allele frequency among worldwide populations not only showed the difference between inland HM people significantly from coastal HM people and other East Asians but also demonstrated the differentiation among different continental populations. The frequency of rs1550623-A located in the intron region of *CDCA7* fixed at 1 in HM and other East Asians and was reported as the strongest risk allele for breast cancer in European (0.8469) [45]. Other risk alleles possessed a low frequency in East Asians but a high frequency in other populations or vice versa (**Figure 8a**). The similar patterns of AFS in some types of cancers also showed comparable genetic basis or pleiotropy at cancer-risk loci. Cancer genetics could help better understand drug discovery and repositioning, stratified screening, informing prevention and informing treatment. Thus, we also analyzed the AFS from 25 phylogenomic loci, which also showed the differentiation pattern from ethnolinguistically diverse populations, suggesting the necessity for genomic testing for drug use, such as warfarin, carbamazepine, clopidogrel, and peginterferon (**Figure 8b**).

**Figure 8.**
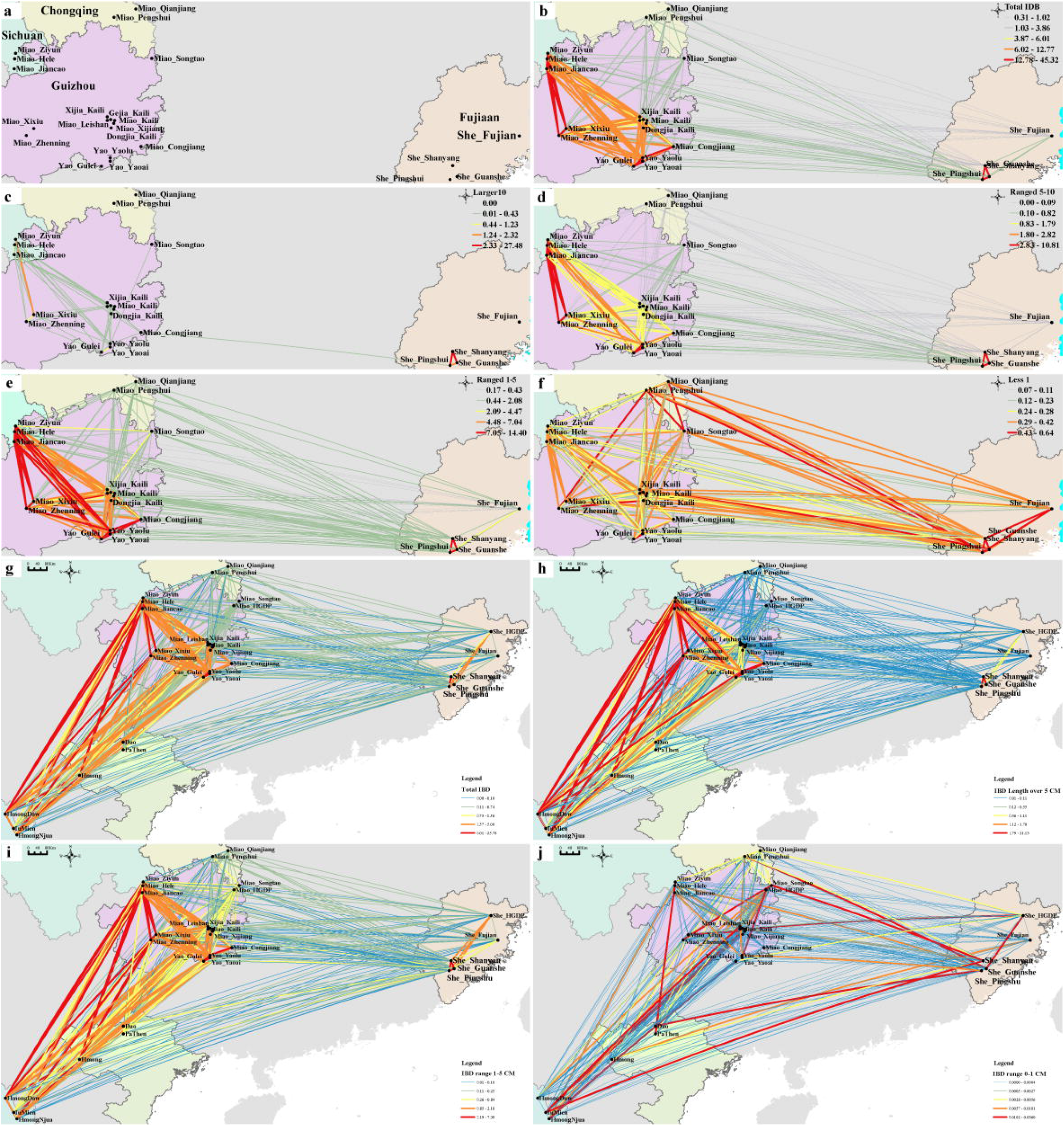
Medical relevance and natural selection signatures among HM-speaking populations. (**a**) Allele frequency spectrum (AFS) of 106 previously reported cancer susceptibility loci among HM people and worldwide reference populations from the NyuWa, GnomAD, and 1KGP. (**b**) The AFS of 25 previously reported pharmacogenomic loci in our dataset and reference groups. (**c**) Allele frequency distribution of natural-selection loci of EDARV370A-rs3827760 A>G among 377 worldwide populations from the 10K_CPGDP, HGDP, and Oceanian genomes. (**d∼e**) Population-specific branch statistics focused on the inland HM people. The top 100 SNPs located in the coding regions were highlighted.

Genetic adaptation of different inland and coastal environments was one of the major evolutionary forces that formed the observed pattern of genetic variations. We used four different statistical methods to search for the signatures of classic sweeps among inland and coastal HM people, including pairwise Fst, integrated haplotype score (iHS), cross-population extended haplotype homozygosity (XPEHH), and population branch statistics (PBS). To explore the shared population-specific adaptation of HM people, we conducted PBS-based genome-wide scanning using the Han_Shaanxi (SXH) and merged European populations from the HGDP as the second and their outgroup reference pulsations, respectively. Several haplotype blocks associated with the immune function and physical traits were evidenced with higher PBS scores (**Figure 8c∼e**). The most robust PBS signature of selection inferred from all three set tests in the form of *PBS*_*HM-SXH-European*_ was *EDAR*, which encoded the soluble ligand ectodysplasin A and a member of the tumor necrosis factor receptor family and regulated the changed tooth morphology and increased scalp hair thickness [46]. Frequency distribution from worldwide populations included in our data, HGDP and 10K_CPGDP (Chinese population genomic diversity project) suggested that East Asian and Native American people harbored a high frequency of the derived allele (**Figure 8c**). Spatiotemporally different ancient DNA evidence from northeastern Asia suggested that *EDAR V370A* likely increased to high frequency after the last glacial maximum [2]. The following identified signals were associated with the immune system and included in *IL6, HLA-DPA1*, and others, which were also the identified highly-differentiated variations (HDV) estimated based on the frequency difference between HM and SXH people. *IGHE*, also named transmembrane protein 121 (*TMEM121*), encoded proteins associated with immunoglobulin receptor binding activity and antigen binding activity was the top HDV. Another important environment-adaptative gene of *SLC24A5* encoded intracellular membrane protein was also identified among coastal and second HM populations. We also conducted PBS analysis in the form PBS_HM1-HM2-SXH_ to further explore the regional HM-specific signals and identified many unique and divergent signals, such as the top identified signatures from the *APCDD1* encoding an inhibitor of the Wnt signaling pathway and *LILRA3* encoding a member of a family of immunoreceptors. We finally used haplotype-based statistical methods to estimate iHS and XPEHH values with the SXH as the reference population, confirming the ancient selection signatures and identifying additional recent or ongoing signals associated with the human immune system diet and exposure to disease.

## Discussion

We presented the complete landscape of population admixture, fine-scale genetic structure and migration history at an unprecedented resolution on the HM population coverage and statistical methods. We merged our generated population data and previously published data genotyped via the Affymetrix Human Origin array and formed the largest and the most comprehensive genomic resource. We collected population data from all different HM ethnicity identities, which included the Miao people from Sichuan, Chongqing, and Guizhou provinces in inland South China, Yao people in the mountainous region of Yungui Plateau, and She people from Fujian province in coastal South China, Dao and Hmong from Vietnam, Hmong people from Thailand, covering over 440 HM people from 34 geographically diverse populations of South China and Mainland Southeast Asia. We comprehensively construct the admixture models using multiple complementary state-of-art statistical and computational methods, including the traditional allele frequency-based PCA, ADMIXTURE, *f*-statistical tests, ALDER-based admixture time estimation, and haplotype-based fine-scale population structure dissection via fineSTRUCTURE, Globetrotter, ChromoPainter and four methods to identify the adaptative signatures. Population effective size and population connection in different historical periods were also estimated via the shared IBD chunk lengths. We additionally explored the paternal and maternal population history via the phylogenetic topology reconstruction, HFS distribution, and Network-based phylogenetic relationship reconstruction. Generally, our results have illuminated the detailed formation process of Chinese and Southeast Asian HM people via the described methods and complex admixture model reconstruction. Fine-scale population structures identified via the sharing alleles and shared IBD lengths were reported at unprecedented resolution. Generally, both autosomal and uniparental genetic evidence revealed the genetic differentiation between inland and coastal southern Chinese HM people and demonstrated the close genetic connection between Chinese coastal HM people and Mainland Southeast Asian HM people, suggesting their recent long-distance migration as documented via the historical records. Medical relevance results of cancer susceptibility genes and pharmacogenomic loci showed that genetic testing in ethnolinguistically diverse populations is important for clinical genome medicine. Recent human local adaptation in HM people has enabled the identification of shared and divergent signatures among geographically different HM people.

### Fine-scale population substructure among HM people and admixture signatures with surrounding neighbors

Large-scale high-coverage WGS projects aiming at East Asians, including the ChinaMAP, NyuWa genomic resources, and other single population admixture history reconstructions focused on Li, Tibetan, Hui and others [5, 6, 47, 48], have provided insights into fine-scale genetic stratification, genetic diversity, dispersal, and admixture landscape. However, HM-speaking populations were a significant lack in the early genomic projects, which are essential for better understanding the genetic diversity of South China and Mainland Southeast Asia. Recent genetic studies from the expanded 1KGP, HGDP, and other Han-based genomic studies have only identified the population differentiation between northern and southern Han Chinese populations [11, 39, 49, 50]. The reported population substructure among geographically Han Chinese was recently confirmed via ancient DNA, illustrating that population stratification between northern Shandong and southern Fujian ancient populations started in the early Neolithic era [15]. Further genomic admixture models further dissected the genetic differences between highland and lowland East Asians or other ethnolinguistically diverse populations, such as Sherpa, Tibetan, and Mongolian [13, 51]. However, only ten She and ten Miao individuals were reported in the HGDP, which showed a close genetic relationship with southern Han Chinese populations. Large-scale population genetic analyses focused on the Chinese populations, included in the 10K_CPGDP or beyond East Asia in the HUGO Pan-Asian SNP project, showed that fine-scale population substructure in the eastern Eurasian strongly associated with the cultural boundaries of geography or language [12, 13, 52], which was also confirmed in our genetic admixture pattern reconstruction among eastern Eurasians. Our previous genetic study focused on the single HM population has identified the previously unidentified genetic cline in Southeast China, which widely existed in the mountainous region of the Yungui Plateau [37, 38]. Liu et al. generated the genome-wide data from Vietnam Hmong, Dao, and PaThen and found that Vietnam HM people received genetic influence from southern East Asians [40]. Kutanan et al. further explored the genetic structure of geographically different Mainland Southeast Asian HM people from three Thailand Hmong populations and found a strong genetic affinity with Vietnam HM [41]. These fragmented HM population data were just reported the basic results based on the sharing alleles and limited to presenting the entire landscape of the patterns of genetic diversity, biological adaptation, medical relevance and population admixture models. Besides, other HM populations from coastal southern Chinese Fujian province are under-representative in early human genetics. Recent human genetic research has emphasized the European bias in genetic studies limited human health equality in non-European populations [9]. The Human Heredity and Health in Africa (H3Africa) Consortium have launched many large-scale population genomic studies to fill the gap in Africa and aimed to present the full spectrum of African genetic diversity. Choudhury et al. recently reported the genomic findings of African migration and healthy based on the genomic data from resources of TryopanoGEN (Trypanosomiasis Genomics Network of the H3Africa Consortium), H3A-Baylor and SAHGP (Southern African Human Genome Programme) [53]. Thus, large-scale population genetic studies from ethnolinguistically diverse Chinese populations are needed to fill the similar gap and provide deep insights into the evolutionary formation of HM people, including their origin, differentiation, and migration past.

We used our collected large-scale genome-wide SNP data of HM people from previously genetically understudied regions and ethnicities and merged it with all modern and ancient East Asians. The estimates of the pairwise genetic distances and patterns of the genetic relationship inferred from the descriptive methods of PCA and ADMIXTURE also confirmed the fine-scale population substructures in the northeast, western, inland southwestern, and coastal southeastern Chinese populations, which was also consistent with the language belongings. Modern population genetic structure patterns were interestingly consistent with the Holocene population structure of ancient East Asians, suggesting the strong population stability of geographically diverse ancient Eastern Eurasians. The ancient genetic structure identified in the Amur River Basin and Mongolian Plateau clustered closely with the Mongolic and Tungusic genetic positions [2]. Similar patterns of genetic stability were observed in Nepal ancients in the Tibetan Plateau [54, 55]. HM genetic cline separated from other southern Chinese populations and formed one unique genetic cline. 500-year-old Gaohuahua people also have a close relationship with modern HM cline [3]. Compared with the inland HM people, coastal Fujian HM people clustered closely to Han Chinese than other HM people. Shared genetic drift and differentiated shared ancestry inferred from *f*_*3*_/*f*_*4*_ statistics in the form of *f*_*3*_(HM, Han; Yoruba) and *f*_*4*_(inland HM, coastal HM; modern and ancient East Asians, Yoruba) confirmed the more genetic influence of Han Chinese and Tai-Kadai people in coastal HM She people. Admixture models based on the qpAdm and qpGraph also suggested more direct gene flow from northern East Asians participating in the formation of the gene pool of HM people. We conducted multiple statistical methods to explore the relationship among them based on the sharing patterns of independent alleles, haplotypes, and uniparental lineages. The genetic relationship between coastal and inland HM people revealed by PCA clustering patterns showed a different affinity between the Yungui Plateau HM people and She people. Formal tests and admixture models via the state-of-art methods of ADMIXTURE, qpGraph, and qpAdm showed that She people had a stronger affinity relationship with Tai-Kadai and Han Chinese populations. We also reconstructed the haplotypes of HM people, dissected the fine-scale population substructure, and differentiated admixture events among different HM populations. We identified a separate branch of the fine-scale structure according to haplotype-based clustering patterns. Paternal lineage reconstruction further revealed the different founding lineages between Fujian and inland HM people.

### Long-distance population movement among HM people and their biological adaptation

Previous modern and ancient genomic evidence has illuminated multiple southward gene flow events that influenced the gene pool of Southeast Asians [40, 41, 56, 57]. The first southward migration may be related to the early Yangtze River rice farmers associated with the AA dispersals. The second and third ones respectively disseminated the AN and TK from South China to Island and Mainland Southeast Asia. The last two were associated with the expansion of HM people from South China and TB people from the Yellow River Basin. Our study aimed to explore the genetic relationship between inland southern Chinese HM people and mainland HM people based on comprehensive admixture modeling, primarily based on sharing reconstructed haplotypes. PCA, ADMIXTURE, and other admixture modeling showed a close genetic relationship between Mainland Southeast Asian HM people and Chinese inland HM people. Here, we additionally reconstructed the phylogeny based on the IBD-based genetic connection. FineSTRUCTURE results confirmed this connection. Finally, shared patterns of different lengths of IBD showed a recent genetic interaction between southwestern Chinese HM and Thailand and Vietnam HM people. Recent genetic studies have identified the long-distance migration in the Eurasian steppe associated with the Mongolian expansion [58]. We also noted that deep coverage of WGS in the next step should be conducted to illuminate more deep insights focused on ethnolinguistically and geographically different HM people, such as the estimates of divergence time and phased genomes for the effective population size estimation and archaic introgression research.

Our newly-generated genomic resource was suitable for illuminating the evolutionary selection force that contributed to the formation of our observed pattern of genetic diversity. We used our sampled populations and four statistical analyses (HDV, PBS, iHS and XPEHH) to identify the genetic variants associated with East Asian living conditions, infectious disease exposure and dietary practices. East Asian-specific adaptative coding variant of the rs3827760 (A>G) within *EDARV370A* has the highest derived allele frequency in HM people. Modeling nonpathological human genetic variation in knockin mice showed that *EDARV370A* controlled the hair thickness and the number of mammary and eccrine glands [46]. Another important human phenotypic diversity is related to skin color. Typical human pigmentation evolution genes of *OCA2, SLC24A5, SLC45A2* and *TYRP1* were evidenced as the light-skinned selected variants in Europeans [46]. African dark-skinned related pigmentation variants were also begun to be explored. Our HM genetic variations also showed the most significant selection signals in *SLC24A5*, one highly penetrant genetic variant of Mendelian disorders and molecularly evidenced in the zebrafish color patterns. Recent WGS-based population analysis focused on northwestern Chinese Hui and Uyghur also identified differenced allele frequency or haplotype homozygosity in the *SLC24A5* gene [47, 59]. The highly significant selection signal of *TMEM121* regulated the skin biopsies in Yunnan minorities [60] and malaria infection-related IL6 variants identified in Hainan Li [48] were identified in our studied HM people. Still, other southern Chinese-dominate selection signatures related to malaria infection (*CR1* and *FREM3*) and fat metabolism-related genes (*FADS*) were not observed, suggesting further high-depth WGS data from more ethnolinguistically diverse populations with larger sample sizes were needed in the Asian genomic projects.

## Conclusions

HM people are essential to the ethnolinguistically diverse South China and Mainland Southeast Asia crossroad regions. It is crucial to understand the entire landscape of genetic diversity, human population evolution, migration, admixture and local adaptation of Asian populations. However, this region is underrepresented in human genomic studies as the European bias in worldwide human genetic research or Han-bias in the Asian cohort studies. To present the global pattern of genetic admixture, adaptative history and fine-scale population genetic structure of ethnolinguistically HM people, we generated the most extensive dataset, including 440 HM people from 34 populations. We conducted population admixture modeling and demographical modeling history reconstruction based on the sharing alleles and haplotypes. We found no genetic traces supported the differentiated genetic structure and geographically close Han and TK people but found elevated northern East Asian ancestry that contributed more to coastal HM people than that to inland ones. Deep population admixture history reconstruction and fine-scale genetic structure dissection illuminated that solid genetic differentiation between inland and coastal HM people supported their independent origin hypothesis. Thus, coastal HM people originated from WuYi mountain, which may be associated with ancient Liangzhu ancestors, and inland one derived from the middle Yangtze River Basin associated with Qujialing, the Shijiahe ancestors. ROH and effective population size estimates and other admixture signatures showed that multiple evolutionary sources constructed to the observed stratification of HM people, such as bottleneck and isolation. We found direct IBD-based genetic evidence supported the association between long-distance population movement and admixture of HM people and the spread of the HM language. Evidence from the adaptative history reconstruction and medical relevance analysis emphasized the shared and divergence patterns of disease susceptibility genetic variations or selected loci. Generally, our results provided deep insights into the formation of HM language and people.

## METHODS AND MATERIALS

### Human subjects

We genotyped 349 HM individuals from 26 ethnolinguistically diverse populations (Miao, Yao, and She) from Sichuan, Chongqing, Guizhou, and Fujian provinces in South China based on the Illumina array platform (**Figure 1a**), where 38 She people from coastal South China was firstly reported here. To present a fully resolved picture of the genetic diversity of HM people, we also collected 20 HM people (10 Miao from Hunan and 10 She from Fujian) from the HGDP [39] and 71 HM individuals (12 Daos, 8 IuMiens, 12 PaThens, and 39 Hmongs) genotyped using the Affymetrix Human Origin array from Vietnam [40] and Thailand [41] from previously investigated populations (personal communication). Our HM dataset included 440 HM people from 34 populations and seven ethnic groups. We also genotyped and reported four AN-speaking Gaoshan people in Fujian to explore the genetic interaction between coastal HM and the AN population. We used PLINK v.1.90 [61] and the King [62] to explore the close relatives within three generations. We estimated the PI_HAT values using the PLINK with the “–genome” parameter. Individual pairs with PI_HAT values larger than 0.15 were further validated the coefficient estimates using King with the parameter of “--related --ibs”.

### Data set and reference populations

To explore the genetic diversity in the context of all East Asian modern and ancient reference populations, we merged our data with the publicly available modern and ancient people from worldwide populations included in the AADR (HO and 1240K datasets) [43] and our previous studies based on the Illumina microarray [13, 20, 21, 37, 38, 63-69]. The high-density dataset was formed by merging all Illumina datasets, also referred to as the Illumina high-density dataset, which included 717,228 SNPs. The high-density dataset was mainly used to conduct the haplotype-based analysis and the uniparental lineage reconstruction. We merged our data with the modern and ancient populations genotyped via the Affymetrix Human Origin array and formed the low-density dataset, which was used to explore the basic pattern of population structure as it included more modern and ancient reference populations. The low-density dataset included 56,814 SNPs. To analyze the complete admixture and interaction landscape between HM people and other ancestral source groups, we merged our dataset with ancient eastern Eurasians included in the 1240K dataset. The Illumina reference populations included two AA Blang and Wa, nine Mongolic-speaking Baoan, Dongxiang, Mongolian, and Yugur, Sinitic-speaking Han and Hui populations from Shaanxi, Sichuan, Gansu, Guizhou, and Fujian provinces, six TB-speaking Pumi, Bai, Hani, Tibetan and Tujia, one Tungusic Manchu, and two Turkic-speaking Kazakh and Salar **(Figure 1d**) [13, 20, 21, 37, 38, 63-69]. The HO modern reference populations included 33 TK people from 26 populations in China and Southeast Asia, 27 Han Chinese people from 6 populations, 276 TB people from 30 Chinese and Southeast Asian populations, 224 AA people from 18 populations, 115 AN people from 13 populations, 30 Japanese and 6 Korean, 140 Mongolic people from 18 populations, 62 Tungusic people from 62 populations [18, 40, 41]. Ancient eastern Eurasians were included in both the HO dataset and the 1240K dataset, which included 48 ancient Yellow River Basin farmers from 20 populations in Shandong, Henan, Shaanxi, and Gansu [15, 18, 70]; 30 ancient people from 13 populations in Amur River Basin or West Liao River Basin [2, 70]; 23 ancient people from 9 population in Guangxi province [3]; 54 ancient people from 9 populations in Fujian province and Taiwan island [15]; 26 ancient people from 10 populations in Japan and Korea Peninsula [17]; 33 ancient people from 7 populations in Nepal in the Tibetan Plateau [54, 71]; 54 ancient people from 9 populations southern Russia around Baikal lake regions [1, 16, 72]; 243 ancient people from 20 populations in the Mongolia Plateau [1]; 18 ancient people from 4 populations in Xinjiang [73].

### Ethics approvement

All included individuals signed the written informed consents and were the unrelated indigenous people in the sampling places. We also provided the necessary genetic counseling and healthy genetic report for the sample donors if they were interested. The study protocol was approved via the medical Ethics committees at North Sichuan Medical College and West China Hospital of Sichuan University.

### Genotyping and quality control

We genotyped 661134 autosomal, 28320 X-chromosomal, 24047 Y-chromosomal, and 3746 mitochondrial SNPs in all Chinese HM people using the Infinium^®^ Global Screening Array. We used PLINK v.1.90 [61] to filter out the variants with missing call rates exceeding 0.05 (--geno: 0.05) and remove samples with missing call rates exceeding 0.1 (--mind: 0.1). Minor allele frequencies (--maf) and Hardy-Weinberg equilibrium tests (--hwe) were also used here. The final HM Illumina dataset included 53,3935 SNPs in 1866 individuals from 123 Chinese populations. The merged HO dataset included 56,814 SNPs from 2139 individuals of 254 populations, and the merged 1240K dataset included the ancient populations that covered 11000 SNPs.

### Global ancestry inference

#### Principal component analysis

We conducted PCA using the smartpca package in the EIGENSOFT 7.2.1 [74] using all modern Chinese populations in the merged Illumina dataset or all modern and ancient eastern Eurasian populations in the merged HO dataset. When ancient people were included in the PCA analysis, we projected ancient populations into the top two coordinates extracted via modern people with the additional parameters (numoutlieriter: 0 and lsqproject: YES). To explore the fine-scale population structure, we subsequently removed modern and ancient people from southern Siberia and northern East Asia. We reran PCA based on the included reference populations from the merged HO dataset. We used R version 3.5.2 and the in-house scripts to plot the scatter diagram.

### Model-based unsupervised ADMIXTURE

We merged our data with different modern and ancient reference populations and ran model-based ADMXUTRE [75] to explore the population substructure among ethnolinguistically diverse or spatiotemporally different East Asians. Using the high-density SNPs in the Illumina dataset, we ran ADMIXTURE by merging our data with Chinese minority groups to avoid the large sample size of Han Chinese populations in the model fitness. To explore the genetic similarities and differences, we also ran ADMIXTURE by merging our data with the HO reference panel. The unsupervised model in the ADMIXTURE version 1.3.0 was used here. We used PLINK v.1.90 [61] and additional parameters (--indep-pairwise 200 25 0.4 and --allow-no-sex) to remove successive variants with more substantial linkage disequilibrium (LD, a squared correlation larger than 0.4) in the 200 SNP sliding windows with an SNP step of 25 SNPs. We ran the admixture models with the predefined ancestral sources (K values) ranging from 2 to 20 and used the cross-validation error estimates to choose the best-fitted models. After pruning the linked SNPs, we used 222,526 SNPs in the Illumina dataset and used 45,725 SNPs in the merged HO datasets.

### Pairwise Fst genetic distances

To evaluate the genetic affinity among different populations in these reference panels, we used PLINK v.1.90 [51] to estimate the Fst genetic distances among any population pair. Pairwise genetic distances were designed with two parameters (--within and --keep-cluster-names).

### Inference of population splits and mixtures

To construct the phylogenetic relationship among these ethnolinguistically diverse populations, we performed phylogenetic history reconstruction using TreeMix v.1.13 [76]. PLINK v.1.90 [61] was used to evaluate the allele frequency in each population, which was used as the input file in the TreeMix-based analysis. We used the parameter of -m to specify the migration edges ranging from 0 to 10 to explore the possible gene flow events. We used the plotting_funcs.R script to visualize each model’s phylogenetic topology and corresponding residual matrix. We have not designed the outgroup population (-root) as we focused on the genetic relationship within East Asians. We used the -k flag (-k 500) to group SNPs to account for the LD. Additional parameters (bootstrap and global) were also used to get the best-fitted model. We also ran Molecular Evolutionary Genetics Analysis (MEGA) [77] based on the Fst genetic matrix to validate the obtained phylogenetic topology, and we obtained the consistent pattern of the major clades.

### Runs of homozygosity (ROH)

We estimated the indicator of genomic autozygosity using PLINK v.1.90 [61] among the high-density Illumina dataset. We set the ROH containing at least 50 SNPs and a total length ≥ 500 kilobases using two parameters (--homozyg-snp 50 and--homozyg-kb 500). Two consecutive SNPs more than 100 kb apart (--homozyg-gap 100) was regarded as independent ROH. The default settings of at least one SNP per 50 kb on average (--homozyg-density 50), the scanning window contains 50 SNPs (--homozyg-window-snp 50), a scanning window hit should contain at most one heterozygous call (--homozyg-window-het 1) and the hit rate of all scanning windows containing the SNP must be at least 0.05 (--homozyg-window-threshold 0.05) were used. We further visualized the ROH distribution of each studied population statistically using R version 3.5.2 via the box plots.

### Shared genetic drift and admixture signature estimation based on allele-frequency-based statistics

To directly measure the genetic affinity within HM people and among HM and other geographically close modern populations, we performed outgroup *f*_*3*_-statistics using the *qp3pop* program in ADMIXTOOLS [43]. As the merged HO dataset included the most comprehensive modern and ancient reference populations, we used *f*_*3*_(HM people, modern Eurasian; Yoruba) to explore the shared genetic affinity between HM people and modern reference populations and used *f*_*3*_(HM people, ancient Eurasian; Yoruba) to measure their genetic relationship with ancient reference populations. We also conducted the three population tests based on the merged Illumina and 1240K datasets. Similarly, we conducted admixture *f*_*3*_-statistics in the form *f*_*3*_(ancestral source1, ancestral source2; HM people) to identify the possible ancestral sources which can produce statistically significant values based on the three datasets. Here, negative *f*_*3*_ values with a Z-score lower than −3 indicated that two sample ancestral sources might be the ancestral source proximities of the targeted populations and also confirmed that the studied population is an admixed population.

### Admixture time estimation based on the decay of linkage disequilibrium

Population admixture can introduce the exponential decay of LD. We used MALDER to test the admixture LD decays and estimate the possible admixture times of HM people [78]. We used multiple modern populations from northern and southern East Asians as the potential ancestral sources and tested all possible source combinations. The exponential curve fitting processes added the minimum distance between two SNP bins (mindis: 0.005 in Morgan) and leave-one-chromosome-out jackknifing (jackknife: YES).

### Genome-wide admixture models based on the *f*_*4*_-statistic tests

We conducted four population tests for targeted HM people based on the individual sample populations and merged populations. We used qpDstat in ADMIXTOOLS [43] to conduct the *f*_*4*_(HM1, HM2; reference populations, Mbuti), *f*_*4*_(studied population1, studied population2; reference populations, Mbuti), and *f*_*4*_(reference populations, studied populations; reference populations, Mbuti). The first form was used to explore the genetic homogeneity and heterogeneity between two included HM populations. The latter two formats were used to test our targeted and reference populations’ differentiated genetic ancestry. We also used the qpWave to confirm the genetic homozygosity between two HM-speaking people and used qpAdm [43] to estimate the admixture proportion with the following outgroups: Mbuti, Russia_Ust_Ishim, Russia_Kostenki14, Papuan, Australia, Mixe, Russia_MA1_HG, Onge, Atayal, and China_Tianyuan. We next used the qpGraph to test the best possible frequency-based admixture models with gene flow events among various alternative models [43].

### Haplotype-based Fine-scale population reconstruction

#### Segmented Haplotype Estimation

More strict filter strategies of missing SNPs and calling rate were performed using PLINK v.1.90 [61] with two parameters (--geno 0.1and --mind 0.1). We used Segmented HAPlotype Estimation & Imputation Tool (shapeit v2.r904) [79] to estimate haplotypes based on the high-density Illumina dataset and modern populations included in the merged HO dataset. Phased haplotypes were estimated with the following parameters to find a good starting point for the estimated haplotypes and get more parsimonious graphs: the number of the burn-in iterations of 10 (--burn 10), the number of iterations of the pruning stage of 10 (--prune 10) and the number of main iterations of 30 (--main 30). We used the model parameters’ default settings and HapMap phase II b37 as the genetic map in the haplotype estimates. The obtained haplotype data was used to explore the fine-scale population admixture via fineSTRUCTURE, identify ancestral proximity and estimate their admixture proportion and time, and screen the natural selection signatures for local adaptation.

### Admixture events inferred from CHROMOPAINTER and fastGLOBETROTTER

To identify ancestral sources, date and describe admixture events of our targeted HM people, we used ChromoPainterv2 [71] to paint the ancestral haplotype composition of our sampled HM populations. We merged our data with 929 lift-over high-coverage whole-genomes from 54 worldwide ethnolinguistically diverse populations and obtained haplotype data using shapeit v2.r904 [79]. Han people from Xi’an (Han_Xi’an), Tibetan_Zase, Daur, Yakut and Hezhen were used as the potential northern East Asian sources of our targets, and Yao_Gulei, Yao_Wangmo, Li_Ledong, Li_Linshui, Li_Wuzhishan and Cambodian were used as the potential southern East Asian sources. We also included other Eurasian populations as the potential ancestral sources in the painting process, including Basque, French, Balochi, Brahui, Pathan and Sindhi. All target HM people were used as the recipients and sources as the donors. The estimated copying vector and painting samples showed the genome-wide patterns of haplotype sharing, which was used to run fastGLOBETROTTER [80] based on the default parameters. We also used sourcefindV2 to confirm the close genetic affinity between our targeted populations and each used ancestral surrogates.

### Painting Chromosomes and fineSTRUCTURE

To dissect the fine-scale population stratifications, we used ChromoPainterv2 [81] to paint all included HM individuals and obtain the co-ancestry matrix. And then, we used fineSTRUCTURE 4.1.0 [81] to dissect the dendrogram based on the haplotype information.

### Identity by descent and effective population size estimation

We used refined-ibd.17Jan20.102.jar [82] to estimate the shared IBD segments within and between HM individuals. We specified 1cM as the minimum length for reported IBD segments (length=0.1). We used the default values of other parameters, including the sliding marker window of 40.0 and the minimum LOD (logarithm of the odds) score for reported IBD segments of 3. We classified IBD fragments into four classes based on the previously published work: short IBD (<1CM), which most likely reflected the ancient genetic connection over 1500 years; intermediate IDB1 (1CM∼5CM) and IBD2 (5∼10CM), which roundly represented the ancient genetic interaction ranging 500∼1500 years ago; and long IBD (>10CM), which was likely the result of recent genetic admixture within 500 years. The population size was used to normalize the average shared IBD, and the IBD matrix was visualized as the heatmap or genetic connection in the map. The software of the ibdne.23Apr20.ae9.jar and the estimated shared IBD fragments were used to estimate the recent effective population size within the recent 50 generations [83].

### Signatures of natural selection

We recognized the natural selection dataset panels based on the estimated genetic affinity and geographical distribution. Here, we separated our dataset into three panels: the coastal panel included the She people, inland panel1 included the unique genetic HM structure, and inland panel2 consisted of Miao people from Chongqing. The identification of natural signatures in inland panel1 was our primary focus. All people showed as outliers or estimated as relatives were removed here. We used PLINK v.1.90 to calculate the pairwise Fst values of each SNP in any population pairs to explore the HDV. And then, we estimated the extent of haplotype homozygosity within the HM population based on the iHS using rehh 2.0 and SelScan [84, 85]. We also used the SelScan to estimate the extent of haplotype homozygosity between HM people and northern and southern Han Chinese populations. PBS was further used to identify the population-specific selection signatures [86].

### Uniparental population history reconstruction

#### Genotyping and quality control

We extracted 24047 Y-chromosomal SNPs and 3746 mitochondrial SNPs from the merged Illumina dataset to explore the paternal and maternal pupation history based on the sharing haplogroups and coalescence processes. We used PLINK v.1.90 to conduct quality control based on the missing SNP rate and missing genotyping rate with two parameters (--geno: 0.1 and --mind: 0.1) [61]. We retained 11369 Y-SNPs in 203 individuals and 1428 mtDNA SNPs in the final quality-control dataset for uniparental evolutionary history reconstruction.

### Haplogroup classification, haplogroup frequency spectrum estimation and clustering analysis

For Y-chromosome haplogroup classification, we used the python package of hGrpr2.py instrumented in HaploGrouper [87] and the LineageTracker [88] to classify the haplogroups. Two additional reference files were used in the HaploGrouper-based analysis, including the treeFileNEW_isogg2019.txt and snpFile_b38_isogg2019.txt. The Chip version was used in the LineageTracker-based analysis. And we also used this software to estimate the haplogroup allele frequency in different levels of the focused terminal lineages. We also performed PCA analysis based on the estimated haplogroup frequency distribution and multi-dimensional scaling analysis (MDS) based on the estimated Fst values from the haplogroup frequency. HaploGrouper and haplogrep were used to classify the maternal haplogroups.

### Phylogeny analysis and Network analysis

We used the BEAST2.0 [89] and the LineageTracker [88] to reconstruct the phylogenetic topology focused on the population divergence, expansion and migration events. BEAUti, Tracer v1.7.2 and FigTree v1.4.4 were used to prepare the intermediate files for BEAST-based analysis and visualize the resulting phylogeny. The BEAST2.0 was also used to reconstruct the maternal phylogeny. Finally, we used the Median Joining Network instructed in the popART [90] to rebuild the network relationship among different haplogroups and populations based on the obtained maternal and paternal genetic variations.

## DATA AVAILABILITY

The Genome-wide variation data were collected from the public dataset of Allen Ancient DNA Resource (AADR) (https://reich.hms.harvard.edu/allen-ancient-dna-resource-aadr-downloadable-genotypes-present-day-and-ancient-dna-data). The used second-analysis results were submitted in the supplementary materials and the raw allele frequency spectrum data are available at ZENODO (https://zenodo.org/).

## ACKNOWLEDGMENTS

This study was supported by the National Natural Science Foundation of China (NSFC 82202078). We thank Prof. Etienne Patin in Institut Pasteur for sharing high-coverage WGS data from Taiwan Island, Island Southeast Asian and Oceanian. We thank Prof. Wibhu Kutanan, Prof. Mark Stoneking and Dr. Dang Liu for sharing genome-wide SNP data from Vietnam, Thailand, and Laos. We also think all volunteer participated to this project.

## AUTHOR CONTRIBUTIONS

G.L.H., M.G.W., C.L. and L.H.W. conceived and designed the study, G.L.H., M.G.W., J.C., Y.L., R.H., P.X.W., S.H.D., Q.X.S., R.K.T., J.B.Y., Z.Y.W., X.F.X., Y.T.S., L.B.Y., L.P.H., J.W.Y. and S.J.N. made all analyses and revised the manuscript. G.L.H. and M.G.W. wrote the manuscript drift. All authers revised the paper.

## CONFLICT OF INTEREST

The author declares no conflict of interest.

## Notes

### Competing Interest Statement

The authors have declared no competing interest.

